# Collective protection against the type VI secretion system in bacteria

**DOI:** 10.1101/2022.09.12.507624

**Authors:** Elisa T. Granato, William P. J. Smith, Kevin R. Foster

**Author notes:** Equal contribution (Elisa T. Granato, William P. J. Smith).

## Abstract

Bacteria commonly face attacks from other strains using the type VI secretion system (T6SS), a molecular speargun that stabs and intoxicates competitors. Here we show how bacteria can work together to collectively defend themselves against these attacks. This project began with an outreach activity: while developing an online computer game of bacterial warfare, we noticed that one strategist (‘Slimy’) that made extracellular polymeric substances (EPS) was able to resist attacks from another strategist that employed the T6SS (‘Stabby’). This observation motivated us to model this scenario more formally, using dedicated agent-based simulations. The model predicts that EPS production can serve as a collective defence mechanism, which protects both producing cells and neighbouring cells that do not make EPS. We then tested our model with a synthetic community that contains a T6SS-wielding attacker (*Acinetobacter baylyi*), and two T6SS-sensitive target strains (*Escherichia coli*) that either secrete EPS, or not. As predicted by our modelling, we find that the production of EPS leads to collective protection against T6SS attacks, where EPS producers protect each other and nearby non-producers. We identify two processes that explain this protection: EPS sharing between cells and a second general mechanism whereby groups of resistant cells shield susceptible cells (‘flank protection’). Our work shows how EPS-producing bacteria can work together to defend themselves from the type VI secretion system.

## INTRODUCTION

Bacteria commonly live in densely-populated, multispecies communities where they must compete for limited space and nutrients (1–3). Life in these environments has driven the evolution of an array of bacterial weapon systems, used to inhibit or kill competing microbes (4, 5). One of the most widespread and sophisticated of these weapons is the type VI secretion system (T6SS), a contractile nanomachine that fires a needle into neighbouring bacteria to deliver toxic effector proteins (6).

Possession of the T6SS can confer a strong ecological advantage to bacteria cells (7–10). Conversely, being on the receiving end of a T6SS attack can be lethal. Defending against the T6SS, however, presents a unique set of problems for targeted cells, as—unlike many other weapon systems—the T6SS does not rely on the target cell’s surface receptors or transport systems. Instead, it uses mechanical force to pierce the target cell’s membrane and deliver its toxic cargo, which limits the potential range of mechanisms that can be employed in defence (11).

In the face of these challenges, here we show how bacteria can work together to protect themselves against T6SS attacks. Individual-level defence mechanisms are known, including immunity proteins, capsule formation, target modification, envelope stress responses, and the removal of susceptibility factors (12–16). However, some of the most powerful defences in bacteria come about when bacteria act as a collective, allowing groups of cells to take advantage of their large numbers (17–22).

This project began when we serendipitously identified a potential collective defence mechanism against the T6SS while developing a video game, as part of an outreach activity. The game—*[Gut Wars](http://www.oum.ox.ac.uk/bacterialworld/gutwars/)*—uses an agent-based model developed by our group for research (17, 23–25) to illustrate different bacterial warfare strategies in the human gut microbiome. Running the game repeatedly, we noticed that ‘Slimy’ strategists, which secrete extracellular polymeric substances (EPS), were particularly resistant against ‘Stabby’ competitors armed with the T6SS. Secretion of EPS is widespread among bacteria and plays a key role in the formation of biofilms (26, 27). From observation of the game, we hypothesized that EPS released from the cell has the potential to protect both the cell that produces it and others around it, thereby functioning as a powerful collective defence mechanism. There is evidence that EPS has the potential to be physically resistant to T6SSs (12, 13, 15). Moreover, biofilm formation—involving EPS secretion—is known to reduce T6SS susceptibility (14), and is a response to bacterial competition via T6SS (28). We therefore decided to investigate whether EPS allows bacteria to work collectively in the face of T6SS attacks.

Here, we develop agent-based models of the competitions between T6SS attackers and EPS-producing target cells. As suggested by the computer game, these analyses predict that secreting EPS is a collective defence mechanism in which producer cells both protect each other, and also cross-protect nearby non-producing cells. We test our predictions using a synthetic community, in which T6SS-wielding *Acinetobacter baylyi* are competing with *Escherichia coli* strains secreting free (i.e. non-membrane-bound) EPS in amounts comparable to clinical *E. coli* isolates. These experiments revealed that EPS can indeed serve as a powerful collective defence against T6SS attacks. We identify two processes by which this collective defence can occur: i) EPS sharing between cells and ii) what we call ‘flank protection’, whereby groups of resistant cells shield non-producers. Interestingly, the latter process appears to be a very general route to collective defence, because it is predicted to occur whatever the underlying mechanism of protection against the T6SS.

## RESULTS

### Agent-based modelling predicts that EPS can protect against T6SS

Bacteria have evolved a wide repertoire of traits and strategies to increase their survival chances in densely-populated communities. We recently devised a simple browser-based computer game—*[Gut Wars](http://www.oum.ox.ac.uk/bacterialworld/gutwars/)*—to showcase some of these competition strategies in a format accessible to children as well as adults. The game asks the user to select different characters representing distinct bacterial competition strategies, which are then pitted against each other in a simulated gut microbiome environment using agent-based modelling (Fig. 1A). Two of the available strategies are EPS production (‘Slimy’) and T6SS firing (‘Stabby’), both common traits in bacteria. While not originally implemented to fulfil this function, we noticed that i) ‘Slimy’ bacteria were surprisingly resistant to T6SS attacks, and ii) the simulated EPS particles covered a large area around the producing cells. This led us to ask whether the secretion of loose (i.e. non-membrane-bound) EPS could protect cells from T6SS firing events, and so help EPS-producing bacteria survive when faced with a T6SS attacker.

**Fig. 1:**
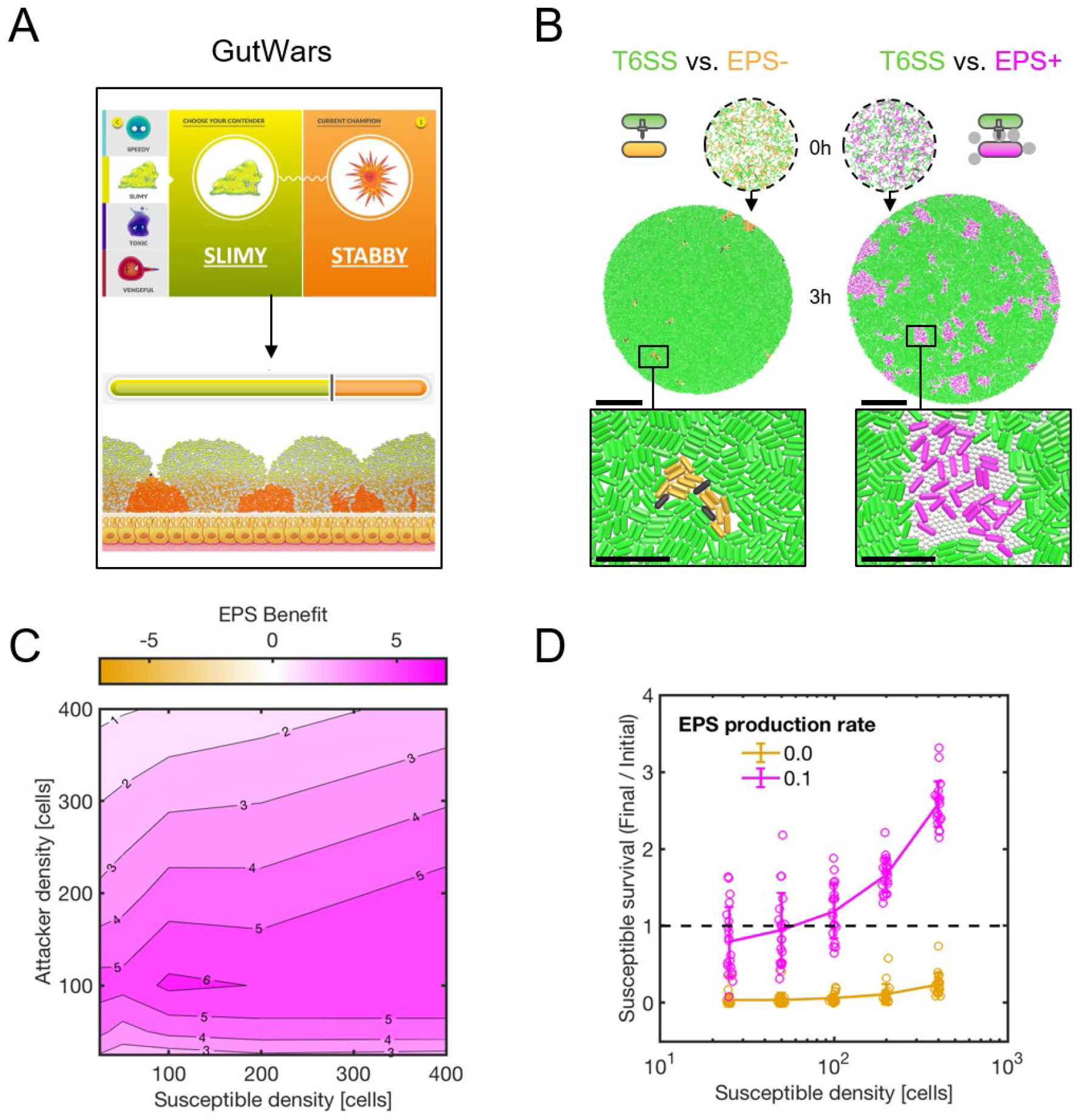
Agent-based modelling of EPS secretion demonstrates collective defence against the T6SS. **(A)** Snapshots of our agent-based video game ‘GutWars’, showing EPS-producing ‘Slimy’ bacteria outcompeting T6SS-armed ‘Stabby’ rivals. **(B)** Agent-based simulations compete T6SS attackers (green) with EPS non-producing (EPS-, P_EPS_ = 0.0) and EPS-producing (EPS+, P_EPS_ = 0.1) susceptible cells (orange and magenta, right and left columns, respectively). Snapshots show i) initial cell seeding (0h), with 400 cells of each type scattered within a circular homeland, and ii) cell configuration after 3 hours’ competition (3h). Scale bars: 50 μm. Zoomed sections highlight dead (black) susceptible cells and distribution of EPS particles. Inset scale bars: 10 μm. **(C)** Contour plot showing measurements of mean EPS benefit (see main text) as a function of initial attacker and susceptible cell density. N=20 simulation replicates per parameter combination. **(D)** Plot showing susceptible final:initial cell counts against initial susceptible density, exemplifying density-dependent benefit of EPS production (fixed attacker density of 400 cells). Individual data points are shown; lines and error bars correspond to data means and standard deviations. Dashed line denotes Final:Initial susceptible ratio = 1, i.e. no net loss or gain of susceptible strain. N=20 simulation replicates per parameter combination; data replotted from (C).

To explore these questions, we implemented EPS secretion within an existing individual-based model of T6SS competition (24, 25). Following previous models (29–31), EPS production is modelled as cells secreting loose, spherical particles into the environment, at a rate *P_EPS_* (see Methods). As they grow, divide, and secrete EPS, producing cells create small pockets of EPS particles around themselves, which helps to separate cells from T6SS-wielding competitors (Fig. 1B). Note, we do not assume here that EPS can physically block T6SS needles, as this effect may not be universal (15). However, were this to occur, the expectation is that it will only increase the protection conferred to cells.

We performed simulation parameter sweeps to identify conditions where EPS production was most protective against T6SS attacks. Comparing simulations with and without EPS (respectively, *P_EPS_* = 0.1, 0), we used a simple metric to quantify the benefit of EPS protection:

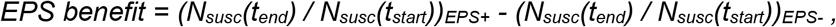

with *N_susc_*(*t*) the number of surviving susceptible cells present at time *t*. Varying the starting number of attacker and susceptible cells shows that EPS production always offers at least some T6SS protection (*EPS benefit* > 0), and that production is most beneficial for intermediate attacker- and susceptible cell densities (Fig. 1C).

We performed additional simulations to test the robustness of EPS-mediated T6SS protection with respect to other simulation parameters (Fig. S1). Repeats of competition simulations with different EPS production rates showed that, as might be expected, susceptible survival decreases as the per-cell rate of EPS secretion is reduced (Fig. S1A); below a minimal secretion rate, survival is indistinguishable from that of a non-producing susceptible strain. We also tested a model in which EPS production is costly, such that susceptible cells suffer a growth penalty proportional to their EPS production rate,

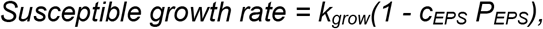

with *k_grow_* the maximum cell growth rate, *P_EPS_* the probability of EPS secretion per simulation timestep, and *C_EPS_* the unit cost of EPS production. For various *C_EPS_* > 0, costly EPS production remains net-beneficial to susceptible cells (Fig. S1B), such that the optimal secretion rate *P_EPS^*^_* > 0. Predictably, EPS investment collapses for excessive costs (Fig. S1B), and is contingent on attacker presence (Fig. S1C). Overall, our simulations predict that production of loose, non-membrane-bound EPS is an effective means for cells to protect themselves against the T6SS, even when costly.

### EPS protection is predicted to function as a collective trait

We next used the simulations to more formally evaluate whether EPS protection is a collective trait. A key feature of many collective traits is a positive relationship between the number of individuals with the trait, and the benefit conferred (32). To look for this, we plotted the benefit of EPS production against the starting number of susceptible (EPS producer) cells, which revealed that the relationship is positive for many levels of EPS secretion (Fig. 1D, S2A, S2B). Note that our metric controls for any density-dependent survival effects emerging solely from changes to cell group size (33). That is, while survival is expected to generally increase with susceptible cell number, this effect becomes much stronger when susceptible cells also produce EPS (Fig. 1D).

Next, we examined the extent to which the benefits from EPS production are shared among cells, which is a second key feature of collective traits. Using the same competition protocol as before, we ran simulations pitting T6SS attackers against susceptible EPS-cells, susceptible EPS+ cells, or both together (Fig. 2A), for a range of initial cell densities. We then quantified the survival of both the EPS+ cells and the EPS-cells when they grew either i) alone with the attacker or ii) paired with the other susceptible cell type (Fig. 2B). As expected when the protective effect of EPS is shared between cells, EPS-survival increased in the presence of EPS+ cells (Fig. 2B, ‘rescue’), and, conversely, EPS+ survival decreased in the presence of EPS-cells (Fig. 2B, ‘exploitation’).

**Fig. 2:**
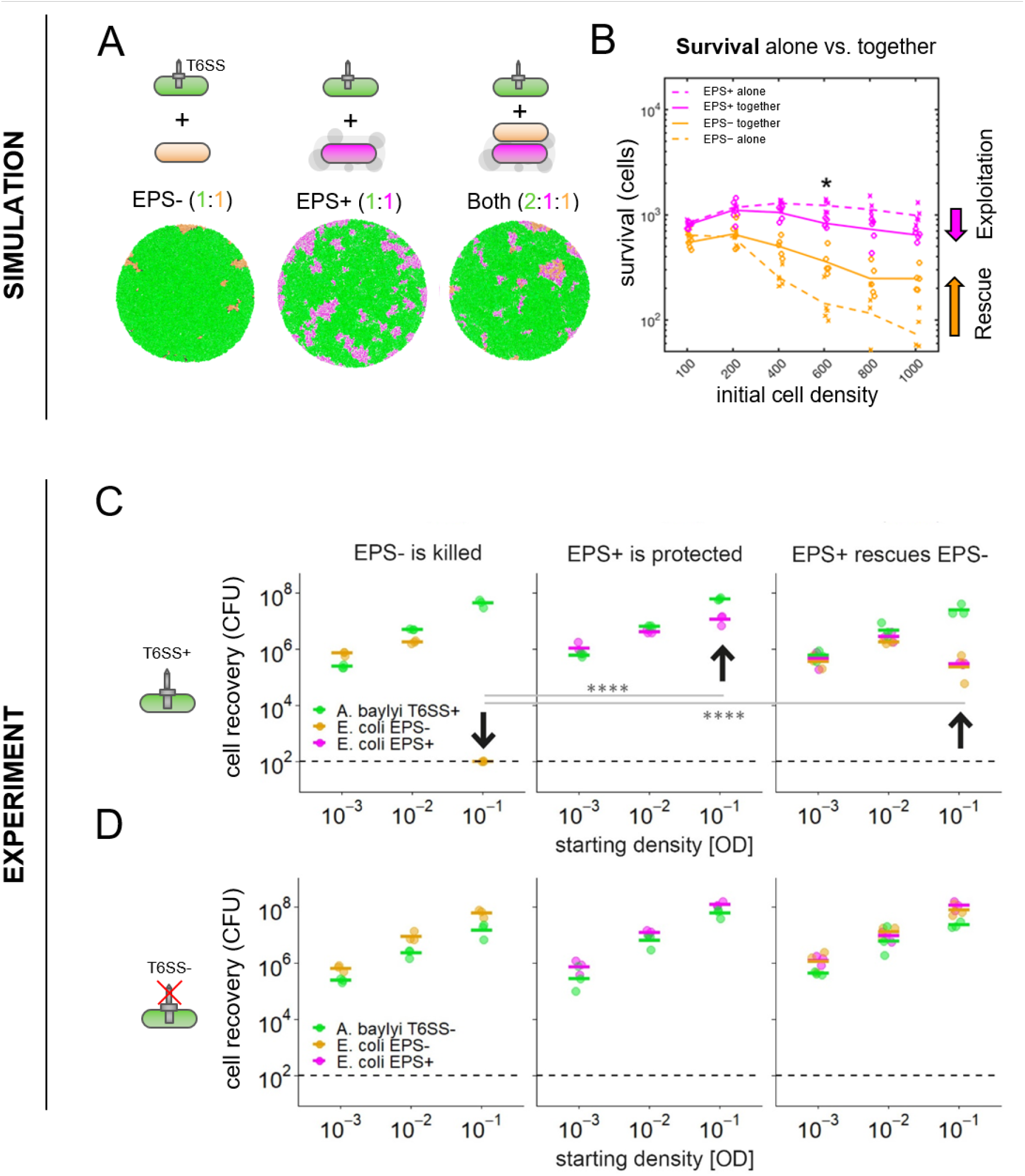
Modelling and experiments show that EPS secretion protects both producer and non-producer *E. coli* from T6SS attacks. **(A)** End-state snapshots of agent-based simulations competing T6SS attackers (green) with EPS- or EPS+ (orange, magenta) strains, or both strains at once. **(B)** Survival of each susceptible strain (EPS+, magenta; EPS-, orange) growing alone against T6SS attackers (dashed lines), or together with the other susceptible strain type (solid lines). Asterisk denotes the scenario depicted in (A). **(C+D)** We conducted analogous competition assays *in vitro* between three bacterial strains: *A. baylyi* ADP1 (T6SS attacker), *E. coli* MG1655 Gen^R^ (EPS-), and *E. coli* MG1655 pRcsA2 (EPS+; colanic acid overproducer). For each strain, cell recovery (CFU) after 6 h of co-culturing at different starting cell densities (optical density; OD) is shown. Individual data points (n=3) and their means are shown as dots and lines, respectively. Arrows highlight key comparisons. **(C)** Survival of *E. coli* EPS- (left), EPS+ (centre) or a 1:1 mix of EPS- and EPS+ (right), when competed against T6SS+ *A. baylyi*. At high seeding density (OD = 10^-1^), EPS+ cells survive pairwise competitions against a T6SS attacker better than EPS-cells (centre and left; p < 0.0001). When co-cultured with EPS+ and competing against the attacker (right), survival of EPS-cells is strongly increased relative to when they face the attacker alone (left; p < 0.0001). When EPS+ cells are co-cultured with EPS- and compete against the attacker (right), EPS+ survival is reduced compared to when they face the attacker alone (left; p < 0.0001). **(D)** Repeat of (C) with *A. baylyi* lacking a functional T6SS. When competed against a T6SS-attacker in pairwise competitions, no difference in cell recovery between EPS+ and EPS-cells was detected (p = 0.66). For both (C) and (D), competitions were analysed per strain combination and seeding density, with statistical analyses on log-transformed data via two-way ANOVA (α = 0.01) with Ŧukey post-hoc tests (****: p < 0.0001).

### A synthetic community demonstrates collective protection via EPS

To test the model’s predictions empirically, we turned to an established experimental system for studying T6SS killing: competitions between a T6SS+ *A. baylyi* attacker strain and a T6SS-susceptible *E. coli* strain (MG1655; (34)). Using their T6SS, *A. baylyi* cells can inject a cocktail of different effector proteins into neighbouring cells which rapidly induce lysis in susceptible *E. coli* (24). To compare the relative T6SS susceptibility of EPS-producing and non-producing *E. coli*, we created *E. coli* MG1655 pRcsA2 (EPS+), a strain capable of undergoing IPTG-inducible production of the exopolysaccharide colanic acid, which is a major component of natural *E. coli* biofilms (35, 36). Colanic acid production is induced via plasmid-based expression of RcsA, the major positive regulator of colanic acid biosynthesis in *E. coli* (37). Its plasmid-free parent MG1655 serves as the non-EPS-producing control strain (EPS-), as MG1655 does not produce significant amounts of colanic acid under the conditions used in this study (Fig. S3).

We first sought to measure the amount of colanic acid secreted by the EPS+ strain, as well as test where it is localised with respect to the cell membrane. Across *E. coli* strains, colanic acid can be secreted both as a capsule tethered to the producing cell membrane, as well as in untethered, loose form (38). The difference between these two states is likely to strongly influence the role of colanic acid in collective protection from external assaults. To benchmark our colanic acid quantification and localization assays, we compared the EPS+ and EPS-strains to six clinical *E. coli* isolates (Table S2). As for the majority of the clinical isolates, phase-contrast microscopy showed that our EPS+ strain has no visible membrane-bound capsule. This observation is consistent with colanic acid being untethered from producing cells and released into the extracellular environment (Fig. S3A). This conclusion was further supported by colanic acid quantification assays where we measured the amount of colanic acid found in total cell extracts vs. supernatant fractions of cultures harvested from agar plates. For the EPS+ strain, the entirety of the detected colanic acid was recovered in the supernatant (Fig. S3B). This same assay also revealed that the amount of loose colanic acid secreted by EPS+ is comparable to that secreted by two clinical isolates (#0172 and #0993, Fig. S3B). The EPS+ *E. coli* strain, therefore, secretes loose colanic acid at rates comparable to naturally occurring *E. coli*, allowing for a direct test of the role of EPS in T6SS-mediated competitions.

Next, we compared the EPS+ and EPS- *E. coli* strains in competition against a T6SS-wielding *A. baylyi* (ADP1; T6SS+) and a T6SS-deficient mutant (ADP1 *ΔtssM;* T6SS-). We seeded a 1:1 mixture of *A. baylyi* with either EPS+ or EPS- *E. coli* at various densities on agar plates and measured strain recovery after 6 h of co-culture using selective plating (Fig. 2C). At low seeding densities, the T6SS is less efficient because it relies on cell-cell-contact for killing (24, 33). At high seeding densities, we expect the T6SS to be maximally effective; this was confirmed in our competition assays, as the EPS-control strain suffered maximal killing at high densities (Fig. 2C, left). In contrast, the EPS+ strain survived much better at high densities (Fig. 2C, centre), with a 10^5^-fold increase in survival against T6SS+ *A. baylyi* compared to EPS-. This advantage was dependent on T6SS-activity, as there was no detectable difference in recovery between EPS+ and EPS-after pairwise competition with the T6SS-*A. baylyi* (Δ*tssM*) mutant (Fig. 2D). To test whether the presence of EPS in the environment alters T6SS activity, we measured firing rates in hundreds of T6SS+ *A. baylyi* cells exposed to cell-free supernatants with and without EPS, and were unable to detect a difference (Fig. S4). These results strongly support our model’s prediction that non-membrane-bound EPS can protect susceptible cells from killing by the T6SS.

We then examined the potential for a producer strain to cross-protect a non-producer, as expected for a collective defence mechanism. We conducted 3-way competitions between T6SS+ *A. baylyi* and both the EPS+ and EPS- *E. coli* (2:1:1 ratio of *A. baylyi*: EPS+: EPS-) and quantified strain recovery as above. At high seeding densities, as predicted by the model, the survival of EPS-increased 1000-fold in the presence of EPS+ (Fig. 2C, right) compared to when they faced the T6SS attacker alone (Fig. 2C, left), indicating that the presence of secreted EPS can rescue otherwise susceptible cells from T6SS attacks even if they do not produce EPS themselves. Further consistent with the modelling, we observed that relative survival of the EPS+ strain decreased in presence of EPS- (Fig. 2C, right) compared to when grown alone with the T6SS attacker (Fig. 2C, left) suggesting that the presence of non-producing cells partially compromises the protective effects of EPS production.

### Cross-protection can emerge from flank protection

Our modelling and experiments show that EPS can function as a collective defence, whose benefit is shared among cells in a colony or biofilm. We next sought to understand the mechanism(s) underpinning this collective property. We initially reasoned that loose EPS would be shared between cells, resulting in the protection of non-producing cells seen in both modelling and experiments. To test this hypothesis, we modified our model so that EPS+ cells no longer secreted loose EPS particles, but instead became intrinsically resistant to T6SS attacks (requiring 2 hits to kill instead of 1), as has been demonstrated in capsule-forming bacteria (15). The goal of this model was to remove the possibility of EPS sharing between cells, so that we could test whether sharing was necessary for the cross-protection of non-producer cells (Fig. 3).

**Fig. 3.**
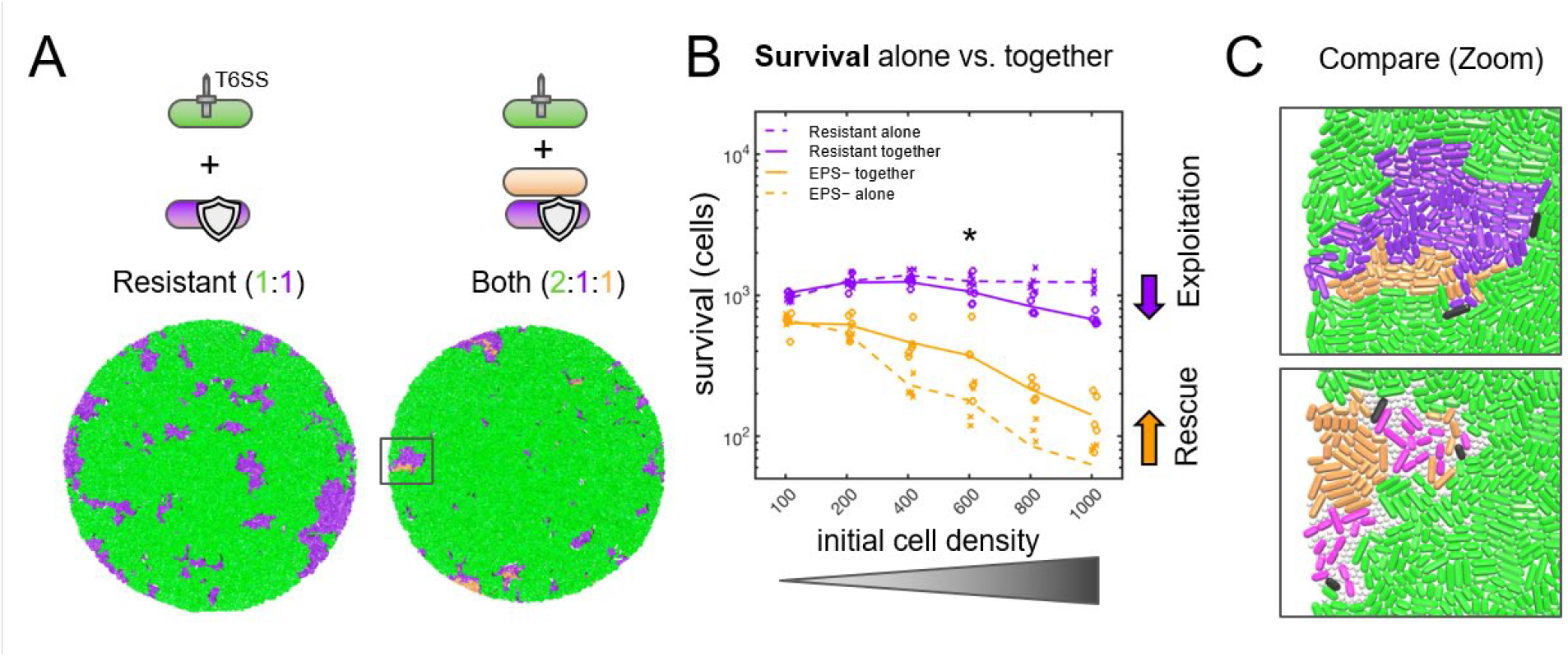
Modelling shows that cross protection can occur without EPS sharing. **(A)** End-state snapshots of agent-based simulations competing T6SS attackers (green) with an EPS-strain (orange), a T6SS-resistant strain (purple) that secretes no EPS particles, or both strains at once. Box corresponds to zoomed section in (C). **(B)** Survival of Resistant strain (purple) and the EPS-susceptible strain (orange) growing alone against T6SS attackers (dashed lines), or together with the other strain type (solid lines). N=5 simulations per parameter combination; lines denote mean survival. **(C)** Magnified views showing Resistant strain shielding EPS-from the attacker (top), similar to when protection occurs via secretion of loose EPS (bottom).

We then competed the resistant cell type against T6SS attackers as before, either in the presence or absence of the EPS-susceptible cell type (Fig. 3A). To our surprise, this again led to collective T6SS defence, which displayed both cross-protection and the same density-dependence seen for loose EPS (Fig. 3B). Close inspection of simulations revealed that resistant cells act as barriers, protecting the flanks of susceptible cell groups by physically separating them from T6SS attackers (Fig. 3C). We therefore term this mechanism ‘flank protection’. Because it involves one set of cells protecting others, flank protection is also a collective effect, but one that can be provided by a purely private resistance mechanism, as well as by loose EPS.

### Time-lapse microscopy reveals both flank protection and EPS sharing

Our modelling raised the possibility that EPS-producing bacteria can collectively defend themselves against T6SS attacks via flank protection, without sharing of EPS between cells. However, an important simplification of the model is that it represents EPS as solid spheres. This limits the ability of EPS to flow past and around cells, restricting sharing in a manner that may not always reflect real EPS (39). We wanted, therefore, to explore both hypotheses—EPS sharing and flank protection—as potential explanations for the collective effects seen in our experiments. To do this, we turned to time-lapse confocal microscopy, which enables direct observation of cells over time (Fig. 4). We competed fluorescent protein-expressing strains of *A. baylyi* (*vipA-sfGFP*, green) with EPS+ *E. coli* (*mScarlet*, magenta) and EPS-*E. coli* (*BFP*, orange) and performed competitions on agar pads as before, except we imaged different locations in the competition area hourly, using confocal microscopy. End-state images of the competition area (Fig. 4A) provided visual confirmation of i) improved survival of EPS+ relative to EPS-, and ii) improved survival of EPS-in the presence of EPS+. Surviving EPS-cells also tended to co-localise with EPS+ cells (bottom right), similar to the behaviour observed in our model.

**Fig. 4:**
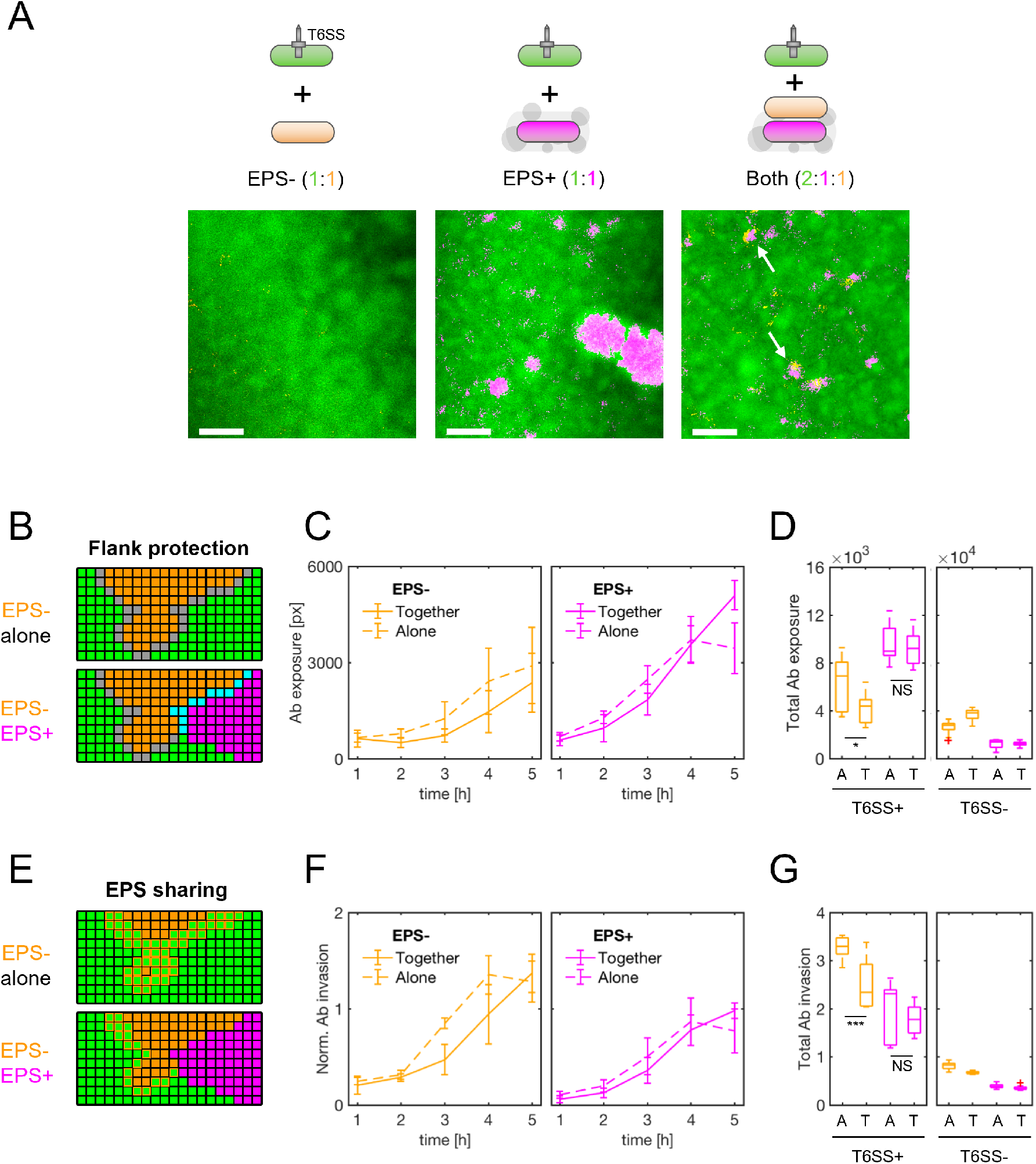
Timelapse microscopy shows that EPS producers cross-protect non-producers via both flank protection and EPS sharing. **(A)** *A. baylyi* attacker (green), EPS- (orange) and EPS+ (magenta) *E. coli* strains expressing fluorescent proteins were mixed and co-cultured as before (Fig. 2C,D), this time analysing competition dynamics using confocal microscopy. Colonies were imaged for 7 h at 1 h intervals, across five 470 × 470 μm frames spanning the colony edge (position 1) to its centre (position 5). Example images shown are zoomed-in areas of interest from position 4 (t = 6 h); scale bars: 50 μm. **(B)** Automated image analysis to identify flank protection: when the EPS-strain is cultured alone (top), its exposure to the attacker (grey boundary) is greater than when the EPS+ strain is occluding part of the EPS-group flank (bottom, cyan boundary). **(C)** Time plots of *A. baylyi* exposure for EPS- (left) and EPS+ (right) strains, grown alone (dashed lines) or together (solid lines), and normalised by initial relative abundance. (**D**) Comparison of total normalised exposure (area under curves in (D)) across all strain pairings, including T6SS-controls. ‘A’ and ‘T’ denote ‘Alone’, ‘Together’. **(E)** Automated image analysis to identify signatures of EPS sharing: when the EPS-strain is cultured alone (top), it is invaded more rapidly by the attacker than when EPS+ is present (bottom), consistent with EPS being shared between *E. coli* strains. **(F)** and **(G):** as with (C, D) but now measuring *A. baylyi* invasion rates, normalised by boundary length. In (C,D,F,G), lines connect data means, and error bars denote standard deviations. Time-lapse data was aggregated over colony positions 2-5; position 1 was excluded for analysis since cells quickly grew beyond a monolayer, confounding our 2-D analysis. N=3 biological replicates; 12 data points per time point. Red crosses denote outlier data points (< Q1 - 1.5 * IQR or > Q3 + 1.5 * IQR). Significance assessed using one-sample Kruskal-Wallis tests: * denotes p = 0.0327; *** denotes p = 0.0004; N.S., non-significant (p > 0.05).

To look for signatures of flank protection and EPS sharing, we used automated image analysis to track the movement of boundaries between *A. baylyi* and *E. coli* cell groups in our time-lapse data. We reasoned that, if cross-protection occurs via flank protection (Fig. 4B), we should observe a reduction in the EPS-strain’s total exposure to *A. baylyi* attackers. More formally, we should observe a reduction in the length of the interface between *A. baylyi* and EPS-cells, when the EPS+ strain is present, versus when it is absent. Alternatively, if cross-protection occurs via the association of loose EPS with EPS-cells (Fig. 4E), then an EPS-cell group should be able to withstand encounters with the T6SS attacker better when EPS+ producers are present as opposed to absent. Note that the tests we develop are not able to estimate the relative importance of flank protection vs EPS sharing, because the measures used in each test are not directly comparable. Nevertheless, we are able to test whether each play a role in the collective benefits of EPS.

#### Flank protection

to test for this, we measured the length of *A. baylyi–EPS-* and *A. baylyi–EPS+* group interfaces over time (see Methods). We compared competitions where the susceptible strains grew alone alongside the attacker with those where both susceptible strains were present. To account for the greater initial abundance of EPS-cells in the former case (2:2 *A. baylyi*: EPS-ratio instead of 2:1:1 *A. baylyi*: EPS-: EPS+), we normalised boundary measurements by the initial relative abundance of each *E. coli* strain (that is, dividing interface lengths by 2 for *E. coli* strains growing alone). As we further discuss in the methods, this normalisation is likely to provide a conservative test of whether flank protection is contributing to cross-protection in experiments (see Methods). Shown in Fig. 4C, these normalised measurements confirm that EPS-indeed has reduced exposure to the attacker when co-cultured with EPS+ (‘together’) vs. without (‘alone’), with the improvement being most pronounced between hours 3 and 4 of competition. Comparing the total attacker exposure (area under curves in Fig. 4C) of each susceptible strain grown alone or together, alongside analogous measurements for T6SS-controls (Fig. 4D), we find that the presence of EPS+ significantly reduces EPS-exposure to *A. baylyi*, consistent with the flank protection hypothesis. Conversely, EPS+ exposure changes only marginally in the presence of EPS-, which is consistent with the inability of the EPS-strain to effectively shield the flanks of EPS+ groups.

#### EPS sharing

to assess this, we measured the rate of *A. baylyi* encroachment into EPS+ and EPS-cell groups (Fig. 4E), normalising this by the total length of interface between *A. baylyi* and each individual susceptible strain (see Methods). The resulting metric, which we term the normalised invasion rate, provides an estimate of how susceptible each *E. coli* strain is to T6SS attack. Crucially, since the normalised invasion rate examines *A. baylyi–EPS+* and *A. baylyi–* EPS-strain interactions separately, it excludes any possible effects of flank protection. Measurements of normalised invasion rates over time showed that the EPS- strain was indeed invaded less rapidly when grown together with the EPS+ strain, vs. when grown alone (Fig. 4F, left). Meanwhile, the EPS+ strain was invaded at a consistently lower rate (Fig. 4F, right) than the EPS- strain, with normalised invasion rates largely unchanged by the presence / absence of the EPS-strain (i.e., no significant exploitation effect detected). Importantly, this variation was contingent on T6SS activity: control experiments showed minimal variation in EPS-strain invasion with EPS+ strain presence when attackers lacked a functional T6SS (Fig. 4G, right).

In conclusion, our analyses suggest that both processes—flank protection and EPS sharing— contribute to the collective benefits of EPS in the face of T6SS attackers.

## DISCUSSION

The production of EPS is a defining property of bacterial biofilms, where the majority of bacteria spend their lives (27). A key challenge for bacteria living in biofilms is the potential for exclusion by competing strains and species (4). Our modelling shows how EPS production can help with this challenge by allowing bacteria to work together and protect themselves against armed competitors. Moreover, an experimental test of our model revealed that EPS dramatically increases *E. coli* survival in competition with T6SS-wielding *A. baylyi*. Here, colanic acid is secreted into the extracellular environment and we find evidence that it has the potential to protect cells other than the producer. Secretion of colanic acid is widespread in *Enterobacteriaceae* such as *E. coli, Klebsiella, Salmonella* and *Enterobacter* (40, 41), where it plays an important role in biofilm formation (35, 36, 42), and confers protection against various environmental stressors such as desiccation, osmotic stress, and oxidative stress (43, 44). Moreover, elevated EPS production (mucoidy) readily evolves in *E. coli* and other species in response to predation by phages (45–47), and membrane-anchored capsules can protect pathogenic bacteria from attacks by the immune system (48–50). EPS-mediated protection from T6SS attacks could therefore also evolve as a by-product of adaptation to other types of stressors, as well as a direct response to T6SS attacks itself.

Both our modelling and experiments showed that EPS non-producing cells are able to survive T6SS attacks better when co-cultured with EPS producers. Our initial hypothesis was that this cross-protection stemmed from the sharing of loose EPS between EPS producers and non-producers. Unexpectedly however, our simulations identified a second cross-protection mechanism, whereby groups of T6SS-resistant producers reduced the exposure of non-producers to attackers (flank protection). We turned to image analysis of our experimental system to examine these possibilities, which suggested that both flank protection and EPS sharing are important in practice. Interestingly, the role we find for EPS secretion implies that secreted EPS can shield non-producing cells, at a range spanning multiple cell lengths, in a manner not seen in the model. However, this finding does not necessarily imply that EPS has evolved *to be shared* with non-producers: it is possible that cross-protection of nearby non-producing cells is a by-product of a diffusible molecule evolved to be shared between cells of the same EPS-producing genotype. Indeed, social evolution theory predicts that such interactions are more likely accidents than evolved altruism (31, 51, 52).

Overall, our findings suggest that EPS production—widespread among bacteria—will frequently result in collective resistance to T6SS attacks, including in cases where EPS is not shared between cells (15). Flank protection in particular has the potential to be highly general: it is predicted to emerge whenever sensitive cell groups interface with resistant cell groups, without assuming anything about the underlying resistance mechanism. Flank protection, therefore, has the potential to be mediated by other mechanisms of T6SS resistance, such as envelope stress responses (13, 16), or the production of immunity proteins (14). Indeed, we hypothesise that flank protection has the potential to occur against any contact-dependent weapon, including against the T4SS, T7SS, and Cdi (contact-dependent growth inhibition) systems (4), adding further complexity to interbacterial interactions. Such effects underline the potential for third parties to weaken interactions between bacterial strains in diverse communities. This idea is known from the study of bacterial cooperation, whereby the presence of other species can limit the potential for cooperating strains to work together (‘social insulation’) (52, 53).

T6SS-wielding bacteria are increasingly being considered as potential biotherapeutics, capable of delivering toxins into pathogenic bacteria (54, 55). Our work cautions that many species have the potential to resist such attacks due to the widespread ability of bacteria to work as collectives. Consistent with this, bacteria are already known to display collective defences against antibiotics (17, 19, 22), predators (56) and phage (20), and biofilm formation continues to be a major issue in industry and medicine (57–60). Strategies that seek to remove problem strains, therefore, needs to consider the ability of bacteria to work both as individuals and as groups.

## MATERIALS AND METHODS

### Agent-based modelling

We ran agent-based model simulations using the open-source modelling software Cellmodeller (61). To simulate T6SS-mediated interactions and EPS particle secretion, we devised additional code modules, available for download at https://github.com/WilliamPJSmith/CellModeller. The key processes incorporated in our model are summarised here.

#### Model description

##### Cell growth and division

Bacterial cells are modelled as exponentially-growing, rigid elastic rods with fixed radius *R*, which push on one another as they elongate and divide (after doubling their initial volume *V_0_*, plus a small random noise term η_div_). All cells are assumed to have equal access to growth-limiting nutrients. Generally, cells pay no growth penalty for using either T6SS needles or EPS secretion, and so a cell’s volume *V* increases over time as *ΔV* = *k_grow_ VΔt*, with *k_grow_* the maximum specific growth rate, and *Δt* the simulation timestep. However, we also explored scenarios in which EPS secretion carried a range of growth costs proportional to the EPS secretion rate (see Fig. S2B,C). Immediately after cell growth, an energy penalty method is applied each timestep to compute cell movements necessary to minimise total cell-cell overlap, subject to viscous drag forces acting on each cell. This process, described previously in detail (23, 61), approximates the elastic repulsion forces acting between cells in physical contact.

##### EPS secretion

Similar to previous work, we model agglomerations of EPS polymers as inert, pseudospherical particles (29–31). EPS+ cells secrete these particles from randomly chosen points on their surface. These particles have radius 0.4 μm, rod segment length 0.01 μm, and are assumed mechanically identical to bacterial cells. The secretion probability per time step, *P_EPS_*, is at most 0.1, such that the max secretion rate is capped at 1 particle per timestep. EPS particles persist indefinitely, and completely ignore the effects of T6SS toxin injection (see below).

##### T6SS interactions

Each time step, T6SS+ cells can fire needles of length *R/2*, projecting orthogonally from randomly chosen sites on their cell surface. The number of firings per cell per unit time is drawn from a Poisson distribution, whose mean is *k_fire_*. After firing, each needle is checked to determine if it comes into contact with any other cell in the population (line-segment method). Subsequent to this contact finding, each cell’s state is updated to indicate the number of T6SS needle hits it sustained. Cells die (cease growing) after being struck *N_hits_* times in this way, and lyse (disappear) following a fixed delay of *1 / k_lysis_*. Each T6SS needle can deliver its toxin payload to at most one cell. T6SS+ cells are assumed completely immune to their own T6SS toxins, such that they effectively ignore all collateral fire from clone-mates. EPS particles do not obstruct T6SS needles, and so EPS particles can only protect bacterial cells from T6SS attacks by enforcing spatial separation of attacker and susceptible cells.

#### Computation and post-processing

All agent-based simulations were run using a 2017 Apple MacBook Pro laptop computer, with concurrent simulations distributed between an Intel 3.1 GHz quadcore i7-7920HQ CPU, an Intel HD 630 Graphics card, and an AMD Radeon Pro 560 Compute Engine. Simulation data were visualised using Paraview software, and analysed using custom Matlab and R scripts.

#### Simulation protocols

All simulations were initialised by randomly scattering different combinations of cells on a flat surface, within a circular ‘homeland’ of radius 50 μm. At simulation time *t* = 0, all cells lie flat on the surface, but have randomised orientations. All simulations were run for 130 time steps (*Δt* = 0.025 h), totalling 3.25 h of simulated time. Depending on conditions (most crucially, the number of EPS particles present), most simulations had a runtime of between 10 and 30 minutes (approximately 10 simulations per hour after concurrency).

### Bacterial growth conditions

All *s*trains were routinely grown overnight in 5 ml LB medium (per L: 10g Tryptone, 10g NaCl, 5g Yeast Extract) in 15 mL polypropylene tubes with agitation (220 rpm). Where necessary, the medium was supplemented with streptomycin (50 μg/mL), ampicillin (100 μg/mL), carbenicillin (100 μg/mL) or gentamycin (15 μg/mL). Routine culturing was carried out on 1.5% w/v LB Agar at 37°C (*E. coli*) or 30°C (*A. baylyi*). All pre-cultures and experiments were carried out at 30°C on 0.8% w/v LB Agar, unless indicated otherwise. Optical density (OD) of liquid cultures was measured at 600 nm. For time-lapse microscopy experiments, samples were kept at 30°C throughout the duration of the experiment using a custom-built incubation chamber.

### Strain construction

For the generation of MG1655 *mScarlet-I*::Tn7 and MG1655 *mTagBFP2*::Tn7, MG1655 cells were transformed with either *pGRG25-Pmax::mScarlet-I* or pGRG25-P*max::mTagBFP2* via electroporation and selected on ampicillin at 30°C. The resulting transformants were cultured for 16 h at 30°C in liquid LB medium supplemented with ampicillin and arabinose (0.5% w/v). A small volume (appr. 5 μL) of these cultures was streaked onto 1.5% w/v LB agar supplemented with arabinose (0.5% w/v) and incubated at 42°C for 12 h. For each construct, three fluorescent colonies were then selected, streaked onto 1.5% w/v LB agar and incubated at 42°C for 12 h. Resulting colonies were tested for ampicillin sensitivity. For the generation of *E. coli* MG1655 *mScarlet-I::Tn7* pRcsA2, MG1655 *mScarlet-I::Tn7* cells were transformed with pRcsA2 (37) via electroporation and selected on 100 μg/mL carbenicillin at 37°C. All plasmids and strains used in this study are listed in Table S2.

### Competition assays

To monitor survival of different *E. coli* strains in competition with *A. baylyi* strains, overnight cultures of *A. baylyi* and *E. coli* were diluted (*A. baylyi* 1:10; *E. coli* 1:100) into 5 mL of fresh LB medium supplemented with appropriate antibiotics (Table S2) and 0.5 mM IPTG to induce production of colanic acid. We used *vipA-sfGFP-labelled A. baylyi* ADP1 as the T6SS-positive competitor, whereas an isogenic *tssM* deletion mutant was used as T6SS-negative control competitor. Cultures were then grown for 3 h with agitation, washed twice with LB medium, and resuspended and normalized in LB medium to an optical density at 600 nm (OD600) of 0.1. An *A. baylyi* strain was then mixed with an *E. coli* strain at a ratio of 1:1 (100 μL: 100 μL), and for each strain combination 5 μL were spotted in triplicates in 24-well plates pre-filled with LB agar supplemented with 0.5 mM IPTG (1 mL per well). For experiments at different seeding densities (Fig. 2), the initial 1:1 mix was further diluted 10-fold (for OD = 0.01) or 100-fold (for OD = 0.001) before being spotted on LB agar. For experiments with three competitors (Fig. 2), one *A. baylyi* strain and two *E. coli* strains were normalized to OD = 0.1, mixed at a ratio of 2:1:1 (*A. baylyi*: *E. coli*: *E. coli*), and further diluted if required before being spotted on LB agar. Plates were then dried in a laminar flow at room temperature for approximately 20 min to allow droplet drying, before being incubated at 30°C for 6 h. After incubation, competition spots were washed off with 500 μL NaCl (0.8%) each and subjected to seven rounds of 10-fold serial dilution. Colony-forming units (CFUs) of *A*. *baylyi* and *E*. *coli* strains were determined by plating on streptomycin (100 μg/mL; selecting for *A. baylyi*), ampicillin/carbenicillin (100 μg/mL; selecting for *E. coli* MG1655 pRcsA2) or gentamycin (15 μg/mL; selecting for *E. coli* MG1655 Gen^R^) media, incubated at 30°C (*A. baylyi*) or 37°C (*E. coli*).

### Determination of T6SS firing rate

To determine T6SS firing rate in *A. baylyi* cells exposed to EPS produced by *E. coli* (Fig. S4A), cells of *E. coli* EPS+ and EPS-were grown overnight for 16 h at 37°C, and 1 mL of bacterial suspension was centrifuged for 5 min at 17000 g to sediment cells. The resulting supernatants were sterile-filtered using 0.2 μm syringe filters (Sartorius), and 5 μL were each pipetted onto separate cut-outs (ca. 5 mm x 5 mm) of 1% w/v LB agarose and left to dry for 20 min at room temperature. In parallel, cells of the focal strain (*A. baylyi* T6SS+ *clpV-mCherry2*) were grown to exponential phase in liquid culture (OD600 ~ 0.8-1.0) before spotting 1 μL of culture onto the agarose pads treated with supernatant, and leaving them to dry for 5 min at room temperature. Pads were then inverted onto a 5-cm diameter glass bottom Petri dish with a 3-cm diameter uncoated n° 1.5 glass window (MatTek Corporation), so that the cells were sandwiched between the agarose and the glass. The sample was then moved to the microscope and imaged immediately. Time-lapse fluorescence microscopy was performed using a Zeiss Axio Observer inverted microscope with a Zeiss Plan-Apochromat 63x oil immersion objective (NA = 1.4) and ZEN Blue software (version 1.1.2.0). Time-lapse images were recorded for 3 minutes at a 10-second interval. Exposure times were 1.05 ms for brightfield and 500 ms for mCherry2 (ex: 589 nm | em: 610 nm). The rate of T6SS firing was measured in a total of n = 916 *A. baylyi* cells via detection of ClpV (*mCherry*) fluorescent foci (24). Each new ClpV focus was assumed to mark one T6SS contraction event in a focal cell. Three separate fields of view at different locations on the same agarose pad were monitored for each treatment. Image analysis was performed using FIJI 2.1.0/1.53c (62). For each field of view, total number of cells and ClpV foci were determined each minute via thresholding, segmentation and object counting in the brightfield channel, and the FIJI built-in ‘Find maxima’ function in the mCherry fluorescence channel, respectively

### Colanic acid quantification

To quantify the amount of colanic acid (CA) produced by different *E. coli* strains (Fig. S2), 1 mL of overnight cultures were washed once with fresh LB medium and density-adjusted to an OD of 1.0. For each strain, 50 μL were then spotted in triplicates in 24-well plates pre-filled with LB agar supplemented with 0.5 mM IPTG (1 mL per well). Plates were then dried in a laminar flow at room temperature for approximately 20 min to allow droplet evaporation, before being incubated at 30°C for 6 h. Cell material was then harvested by adding 1 mL of sterile ddH_2_O to the well and pipetting up and down until all material was suspended, and then transferred to a sterile 1.5 mL tube. Tubes were vortexed for 5-7 seconds, and the OD600 was determined for all replicates. Of these volumes, twice 400 μL were then transferred to two separate reaction tubes pre-filled with 600 μL ddH_2_O: one for quantification of total CA, and one for quantification of CA secreted into the extracellular environment. For quantification of total CA (Fig. S3), samples were boiled for 20 min at 95°C, centrifuged at 17’000 g for 20 min, and 1 mL of supernatant was used for further processing. For quantification of CA in the extracellular environment (Fig. S3), samples were centrifuged at 17’000 g for 20 min, and 1 mL of supernatant was used for further processing. To test whether sterile-filtering removes EPS from supernatants (Fig. S4B), *E. coli* MG1655 pRcsA2 (EPS+) and MG1655 Gen^R^ (EPS-) strains were grown overnight for 16 h at 37°C, and 1 mL of bacterial suspension was centrifuged for 5 min at 17000 g to sediment cells. The resulting supernatants were either sterile-filtered using 0.2 μm syringe filters (Sartorius) or left unfiltered, and 900uL of supernatant was used for further processing. Relative CA production was then determined through the quantification of L-Fucose (63). Briefly, samples were boiled with 4.5 mL H_2_SO_4_·H_2_O (6:1 H_2_SO_4_ and H_2_O) for 20 min and cooled to room temperature. Then, 900 μL of each sample was transferred to a spectrophotometer cuvette and absorbance was measured at 396 nm (A_396_) and 427 nm (A_427_). To each cuvette was then added 100 μL of freshly prepared L-cysteine hydrochloride (1M), mixed by pipetting, and absorbance was again measured at 396 nm (A-cy_396_) and 427 nm (A-cy_427_). The concentration of L-Fucose was then determined by calculating (A-cy_396_ – A_396_) – (A-cy_427_ – (A_427_) and applying a previously generated L-Fucose standard curve (0-100 μg/m?).

### Confocal microscopy

To image competition dynamics, fluorescently tagged versions of all relevant strains (Table S2) were pre-cultured and density-adjusted to OD600 = 0.1 as described in ‘Competition assays’. For each strain combination, 5 μL were then spotted onto a glass bottom Petri dish filled with 8 mL 0.8% w/v LB agar supplemented with 0.5 mM IPTG. After drying for ten minutes at room temperature, the plate was inverted to allow for immediate imaging of the competition area through air. Time-lapse confocal imaging was then carried out using a Zeiss Plan-Apochromat 10x objective (NA = 0.45) on a Zeiss LSM880 confocal laser scanning unit using ZEN Black software (version 14.0.18.201). Images (each covering an area of 472.33 μm × 472.33 μm) were acquired hourly for 7 h, over 5 different colony positions, spanning in total the entire radius of the competition area (edge to centre). Three biological replicates were carried out for each strain combination.

### Confocal image analysis

#### Image segmentation and stabilization

Following acquisition, raw confocal image data were analyzed using FIJI (62). Multichannel images were split into individual channels, with each channel subjected to separate background subtraction (rolling ball radius 50 px) and conversion to binary image data via user-defined thresholds. To maximise the accuracy of cell group boundary tracking, we stabilised images using the FIJI ‘StackReg’ plugin (64), applied to the GFP (*A. baylyi*) channel. This stabilisation made it necessary to exclude later time points (t = 6,7 h) as these were typically confluent, confounding accurate registration of images. We also excluded colony position 1 (colony edge); here, coffee-ring effects (65) consistently led to the formation of 3-D cell groups as early as 2 h into the competition. As our image analysis is limited to 2-D, we were unable to accurately track inter-strain boundaries at this position. For visualization purposes, mtagBFP2 fluorescence is false-coloured orange in all images throughout the manuscript.

#### Inter-strain boundary length (Flank protection)

Beginning with our stabilised, segmented timelapse images, we measured the lengths of *A. baylyi*–EPS-group interfaces by dilating *A. baylyi* cell group images (MATLAB *imdilate* method; 3 pixel spherical structuring element), computing their overlap (in pixels) with EPS+ cell groups, and then summing overlapping pixels. We verified that performing this procedure in reverse (dilate EPS-onto *A. baylyi*) yielded similar boundary pixel counts. To account for the greater initial abundance of EPS-cells when EPS+ was absent (2:2 *A. baylyi*: EPS-ratio instead of 2:1:1 *A. baylyi*: EPS-: EPS+), we normalised these boundary measurements by the initial relative abundance of the EPS-strain (that is, dividing interface lengths by 2 for *E. coli* strains growing alone). EPS+ exposure to the attacker was calculated and normalised in the same way.

This normalisation likely underestimates the role played by flank protection, because it assumes that a focal strain growing alone with the attacker will start with double the exposure as compared to when it is growing with the other *E. coli* genotype. In reality, the exposure in the single *E. coli* treatments is likely to be less than double, so that the normalisation leads to a lower estimated exposure than reality. To understand why, it is helpful to consider what happens when additional *E. coli* cells are present at the beginning of an experiment. In some cases, these will form independent microcolonies that increase the interface with *A. baylyi*, as our normalisation reflects. However, in other cases these additional *E. coli* cells will simply merge with existing groups instead of growing alone. As a result, starting with more cells will not always proportionally increase attacker exposure. For these reasons, we may underestimate the exposure of *E. coli* growing alone with the attacker. Nevertheless, our measure provides a conservative test of whether flank protection is contributing to cross-protection in experiments, because we still see that exposure of EPS-is reduced by the presence of EPS+.

#### Normalised invasion rates (EPS sharing)

To estimate T6SS killing rates at inter-strain boundaries (Fig. 4), we tracked changes in strain occupancy at each pixel across consecutive time-lapse images, using custom MATLAB scripts. We compared consecutive time points (t-1, t; t=1,2,3,4,5 h) in each stabilised time lapse. Any pixel where EPS-channel signal was lost, and replaced with new *A. baylyi* signal, was marked as being ‘invaded’ by *A. baylyi*, giving a measure of EPS-cell killing by the attacker. For each time point t, EPS-invasion pixel counts were normalised by the total amount of *A. baylyi*–EPS-contact found in the t-1 image. EPS+ invasion was calculated and normalised in the same way.

### Statistical Analysis

Statistical analysis and data visualization were performed using RStudio version 1.2.5033 (66) and packages dplyr (67), Rmisc (68), ggplot2 (69), and cowplot (70); as well as Matlab version 2017b (9.3.0.713579). For all statistical tests, the significance level α was set to 0.05. To test whether cell recovery differed between competitions (Fig. 2), two-way ANOVA of log-transformed CFU values with a post-hoc Tukey’s HSD test was performed. To test whether colanic acid production differed between strains (Fig. S3B), and whether firing rate differed when cells were exposed to EPS (Fig. S4A), a two-sided Wilcoxon rank sum test was performed. To test whether filtering removed EPS from supernatants (Fig. S4B), a two-sided, paired samples Wilcoxon signed rank test was performed. To test whether total attacker exposure or total normalised invasion of the EPS-strain changed when grown alongside the EPS+ strain (or vice versa; Figs. 4D and 4G), we used one-sample Kruskal-Wallis tests. Statistical details (sample size, test values, p-values) for each experiment are provided in the respective figure legends.

## ACKNOWLEDGEMENTS

We thank Marek Basler, Colin Kleanthous and Frances Spragge for strains, Chris French and Thomas Meiller-Legrand for plasmids, and Erik Bakkeren, Ravinash Krishna Kumar, Jake Palmer, and Rachel Wheatley for comments on the manuscript. We are indebted to Scott Billings and the Oxford Natural History Museum staff, with whom we developed the *GutWars* video game exhibit, for inspiring this project. ETG is funded by a Postdoc Mobility Fellowship from the Swiss National Science Foundation (P400PB_183878) and a BBSRC Discovery Fellowship (BB/V004328/1). WPJS and KRF are supported by the National Institutes of Health (project numbers R01AI093771 and R01AI120633), by European Research Council Grant 787932, and by Wellcome Trust Investigator award 209397/Z/17/Z. WPJS is also funded by a Sir Henry Wellcome Postdoctoral fellowship award, 222795/Z/21/Z.

## Notes

### Competing Interest Statement

The authors have declared no competing interest.

### Summary of Updates

Edits to main manuscript text. Updated Fig. 4. Added additional analyses regarding 'flank protection' in results section.

