## Supplementary Material for "Collective protection against the type VI secretion system in bacteria"

*Equal contribution

#### SUPPLEMENTARY FIGURES

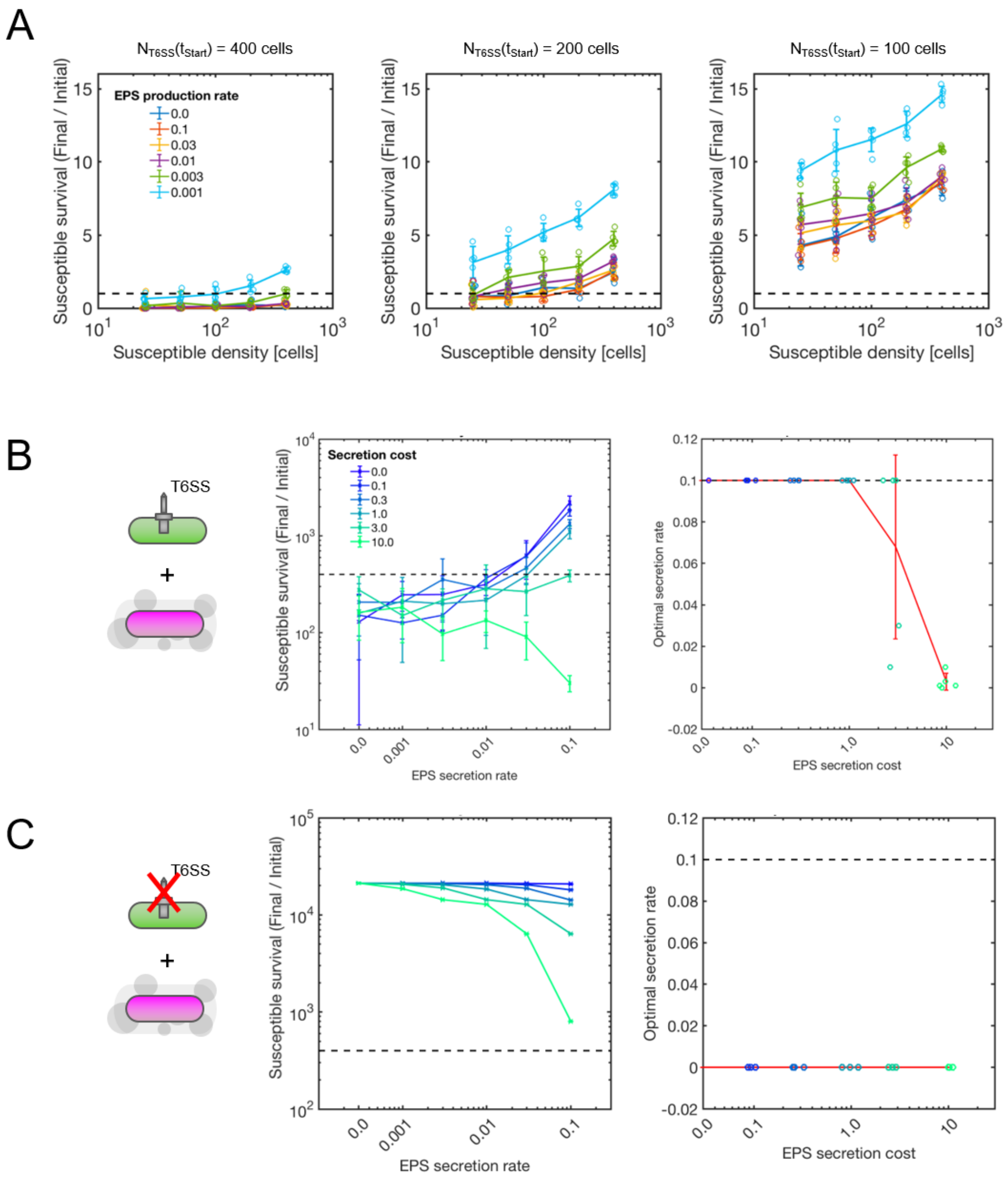


**Fig S1: The benefits of EPS protection vary with EPS secretion rate, cost and attacker density. (A)** Measurements of final:initial susceptible cell counts as a function of initial susceptible cell density, for increasing EPS secretion rates $P_{EPS}$ (see shared legend). From left to right, panels show data for three different starting attacker densities ($N_{T6SS}(t_{Start})$ = 400, 200 and 100). Individual data points are shown; lines and error bars indicate data means and standard deviations, respectively. N = 5 simulation replicates per case. **(B)** Measurements of final:initial susceptible cell counts (left) against T6SS+ attackers ($k_{fire}$: 200.0 firings cell$^{-1}$ h$^{-1}$), as a function of EPS secretion rate $P_{EPS}$, for increasing EPS cost values $c_{EPS}$ (see legend). Each simulation begins with 400 attacker and 400 susceptible cells. Lines and error bars respectively show data means and standard deviations; N = 5 replicates per case. Right: plots of optimal secretion rate, showing the $P_{EPS}$ values that maximised susceptible cell survival as a function of EPS cost $c_{EPS}$. Note that $P_{EPS}$ = 0.1 is the maximum EPS secretion rate we allow, and so true optima may exceed those shown here. Data points are colorcoded according to the legend shown in the left panel; red lines and bars indicate mean and standard deviation of measured optima. **(C)** Analogous to **(B)**, except with susceptibles competing against a T6SS- attacker ($k_{fire}$ = 0.0 firings cell$^{-1}$ h$^{-1}$).

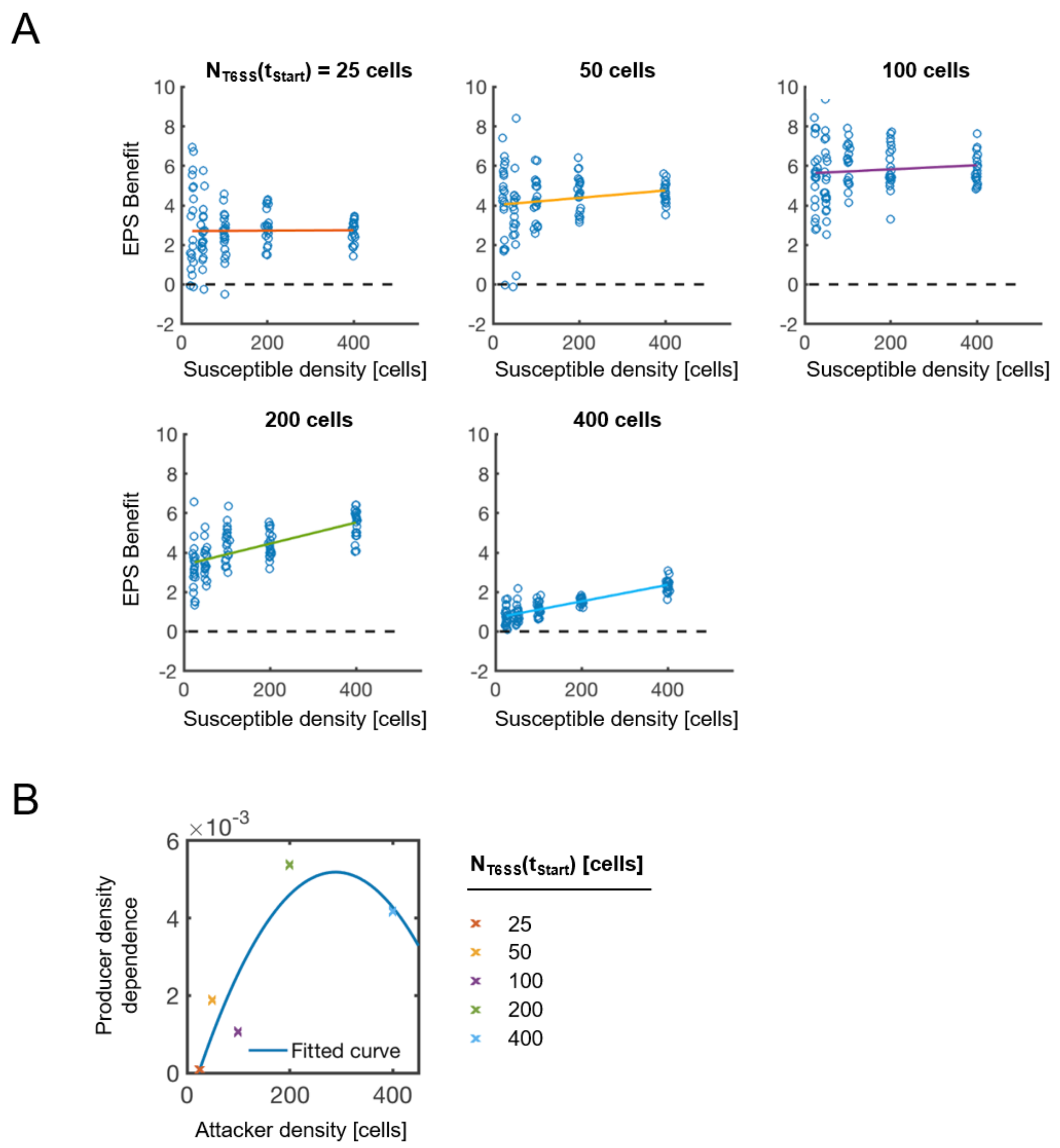


**Fig. S2: EPS production often confers a density-dependent survival benefit against T6SS attacks.** (**A**) Plots of EPS Benefit (see main text) against initial susceptible cell density for increasing initial attacker densities ($N_{T6SS}(t_{Start})$ values indicated above panels). Individual data points shown alongside lines-of-best-fit, fitted using linear regression. Dashed lines correspond to EPS Benefit = 0. N = 20 simulation replicates per parameter combination. (**B**) Summary plot showing gradients of fitted lines in (A) ("Producer density dependence") as a function of attacker cell density. Individual data points are shown alongside a fitted quadratic curve (least-squares fitting, $R^2$ = 0.5588).

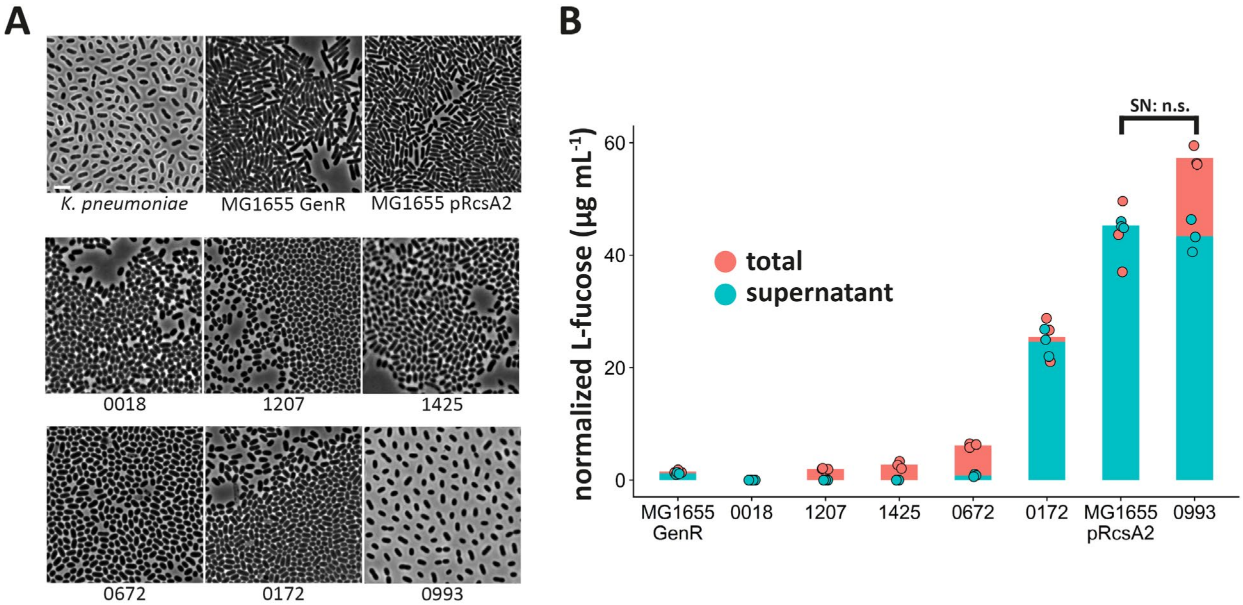


**Fig. S3. EPS phenotype of engineered *E. coli* EPS-producing strain is comparable to that of clinical *E. coli* isolates. (A)** Representative phase-contrast micrographs of EPS non-producing (MG1655 Gen$^R$) and EPS-producing (MG1655 pRcsA2) *E. coli* used in this study, as well as of several clinical *E. coli* isolates (see Table S2). Strains expressing membrane-bound capsules (e.g. isolate 0993) display large gaps between cells even if seeded at high densities. *Klebsiella pneumoniae* is shown as an example of a typical membrane-bound capsule phenotype. Scale bar, 5 µm. **(B)** To estimate relative colanic acid production, monocultures of different *E. coli* strains grown on plates were subjected to a spectrophotometric assay for L-fucose. Individual dots represent L-fucose concentrations normalized to optical density for total cell extracts (red) and supernatants (turquoise), respectively. Coloured bar plots represent mean values of three biological replicates. For each biological replicate, we measured both total cell extracts and supernatants. To test for differences in L-fucose concentration in supernatants (SN), we used a two-sided Wilcoxon rank sum test (W = 6, p = 0.7; n.s., not significant).

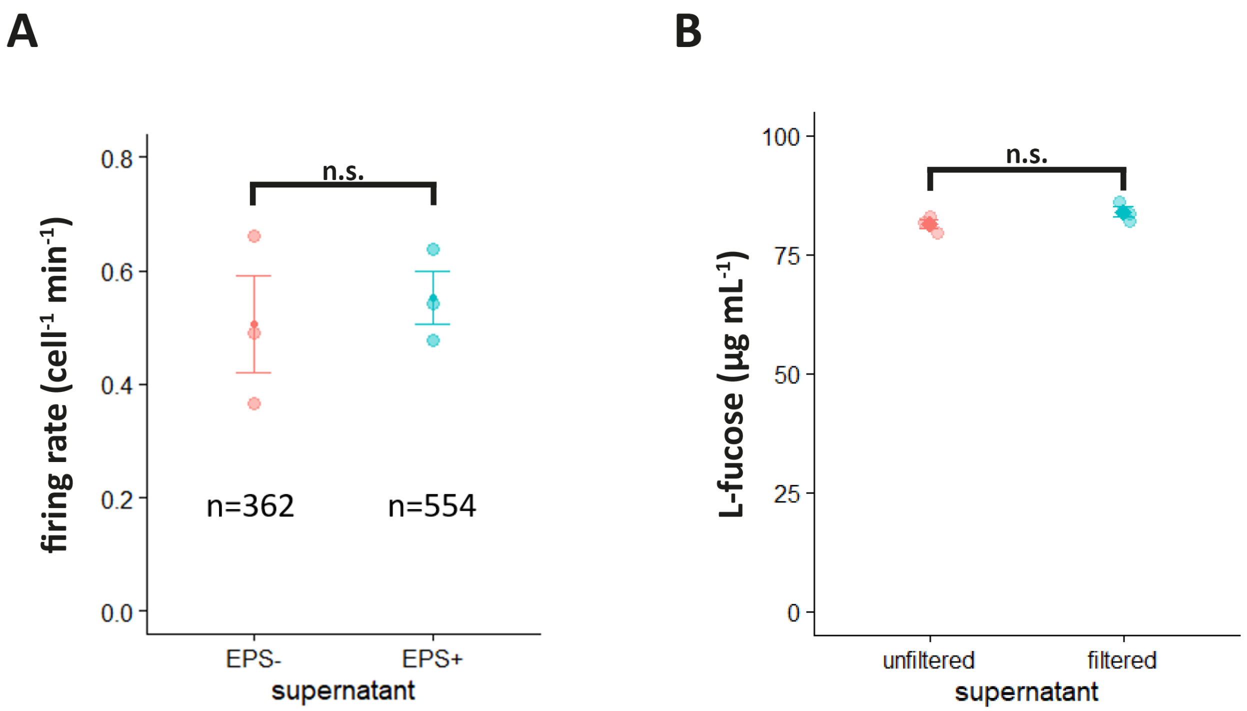


**Fig S4: T6SS firing rate is unaffected by the presence of exopolysaccharides. (A)** T6SS firing rates in *A. baylyi* T6SS+ (*clpV-mcherry2*) exposed to cell-free supernatants of EPS- and EPS+ *E. coli*. Firing rates were determined by time-lapse microscopy and monitoring the appearance of fluorescent ClpV foci in focal cells over time. Darker shade dots represent mean values across three replicates. Error bars denote standard error of the mean. To test for differences in firing rate, we used a two-sided Wilcoxon rank sum test ($W = 4$, $p = 1$; n.s., not significant). n = total number of cells observed for each treatment. **(B)** To test whether sterile-filtering removes EPS from culture supernatants, filtered and unfiltered supernatants of MG1655 pRcsA2 monocultures (n=3) were subjected to a spectrophotometric assay for L-fucose. Individual dots represent L-fucose concentrations for either unfiltered supernatants, or supernatants filtered through a 0.2 µm filter. Darker shade dots represent mean values of three biological replicates. Error bars denote standard error of the mean. For each biological replicate, we measured both filtered and unfiltered supernatant. To test for differences in L-fucose concentration, we used a two-sided, paired samples Wilcoxon signed rank test ($V = 0$, $p = 0.25$; n.s., not significant).

### SUPPLEMENTARY TABLES

#### Table S1. Model parameters used in this study

| TYPE | Parameter | Symbol | Value(s) | Units | Source |
|---|---|---|---|---|---|
| T6SS attacks | T6SS firing rate | $k_{fire}$ | 200.0 | firings cell$^{-1}$h$^{-1}$ | This study |
| | Lysis delay | $1 / k_{lysis}$ | 0.125 | h | (1) |
| | Lethal hit threshold | $N_{hits}$ | 1, Infinity | - | Estimated from (2) |
| | Extracellular needle length | $L_{needle}$ | 0.5 | microns | (1) |
| | Min. needle penetration | $L_{penetration}$ | 0.01 | microns | (1) |
| Cells | Max. specific growth rate | $k_{grow}$ | 1.0 | h$^{-1}$ | (3) |
| | Cell radius | $R$ | 0.5 | microns | Estimated from (2) |
| | Cell volume at birth | $V_0$ | 1.16 | microns$^3$ | (4) |
| | Cell division volume noise | $\eta_{div}$ | 9 | % | (4) |
| | Cell division orientation noise | $\eta_{orient}$ | 0.2 | % | (4) |
| EPS | EPS particle radius | $R_{EPS}$ | 0.4 | microns | This study |
| | EPS particle segment length | $L_{EPS}$ | 0.01 | microns | This study |
| | EPS secretion probability per timestep | $P_{EPS}$ | 0.0-0.1 | - | This study |
| | EPS secretion cost | $c_{EPS}$ | 0.0-10.0 | - | This study |
| Numerical | Simulation timestep | $\Delta t$ | 0.025 | h | (3) |
| | Cell / needle sorting grid size | $h$ | 10 | microns | (1) |
| | Conjugate gradient absolute tolerance | $e_{CG}$ | 0.001 | - | (3) |
| | Max. contact iterations | $Max_{iter}$ | 8 | - | (3) |
| | Regularization weight | $\alpha$ | 0.04 | - | (4) |
| | Growth restriction factor | $1/\gamma$ | 0.002 | - | (4) |

**Table S2. Strains and plasmids used in this study**

| STRAINS | Name | Species | Genotype | Isolation Site | Source |
|---|---|---|---|---|---|
| | T6SS+ | *Acinetobacter baylyi* | ADP1 (vipA)-sfGFP; $Str^R$ | - | (1) |
| | T6SS- | *Acinetobacter baylyi* | ADP1 ΔtssM (vipA)-sfGFP; $Str^R$ | - | Marek Basler |
| | T6SS+ clpV-mCherry2 | *Acinetobacter baylyi* | ADP1 (clpV)-mCherry2; $Str^R$ | - | Marek Basler |
| | MG1655 | *Escherichia coli* | F- lambda- ilvG- rfb-50 rph-1 | - | Colin Kleanthous |
| | EPS- | *Escherichia coli* | MG1655 $Gen^R$ | - | (5) |
| | EPS- (BFP) | *Escherichia coli* | MG1655 mTagBFP2::Tn7 | - | This study |
| | EPS+ | *Escherichia coli* | MG1655 pRcsA2; $Amp^R$ | - | This study |
| | EPS+ (mScarlet) | *Escherichia coli* | MG1655 mScarlet-I::Tn7 pRcsA2; $Amp^R$ | - | This study |
| | #0018 | *Escherichia coli* | unsequenced clinical isolate #19Y000018 | Urine | Nottingham Pathogen Bank |
| | #1207 | *Escherichia coli* | unsequenced clinical isolate #17Y001207 | Pus (Appendix) | Nottingham Pathogen Bank |
| | #1425 | *Escherichia coli* | unsequenced clinical isolate #18Y001425 | Fluid (Pancreas) | Nottingham Pathogen Bank |
| | #0672 | *Escherichia coli* | unsequenced clinical isolate #17Y000672 | Pus (Appendix) | Nottingham Pathogen Bank |
| | #0172 | *Escherichia coli* | unsequenced clinical isolate #18Y000172 | Urine | Nottingham Pathogen Bank |
| | #0993 | *Escherichia coli* | unsequenced clinical isolate #18Y000993 | Fluid (Pancreas) | Nottingham Pathogen Bank |
| | *K. pneumoniae* | *Klebsiella pneumoniae* subsp. *pneumoniae* DSM 30104 | - | - | Frances Spragge |

| PLASMIDS | Name | Description | |
|---|---|---|---|
| | pGRG25-*Pmax*::mScarlet-I | pGRG25 vector for Tn7 integration (6), expressing mScarlet-I from the Pmax promoter (7). Amp$^R$ | Thomas Meiller-Legrand |
| | pGRG25-*Pmax*::mTagBFP2 | pGRG25 vector for Tn7 integration (6), expressing mTagBFP2 from the Pmax promoter (7). Amp$^R$ | Thomas Meiller-Legrand |
| | pRcsA2 | Encodes rcsA under the control of the lac promoter. Allows IPTG-inducible production of RcsA, which upregulates the expression of colanic acid synthesis genes from the chromosome. Amp$^R$ | (8) |
